# A novel sample-delivery method for powder X-ray diffraction at Turkish Light Source

**DOI:** 10.64898/2026.04.09.717569

**Authors:** Esra Ayan, Abdullah Kepceoglu, Arif Mermer

## Abstract

Powder X-ray diffraction (PXRD) measurements performed on platforms originally designed for single-crystal diffraction are strongly affected by how the powder sample is presented to the X-ray beam, including the delivery configuration and support geometry. Here, we developed a modified Terasaki-plate-based sample-delivery method for PXRD using a laboratory single-crystal diffractometer implemented with the XtalCheck-S plate-reader operational mode at Turkish Light Source. The method was regarded under comparable measurement conditions relative to a standard loop/pin-based and a grease-based Terasaki setup using 5-{[4-(2-Methoxyphenyl)piperazin-1-yl]methyl}-4-ethyl-4H-1,2,4-triazole-3-thiol as a model analyte. The loop-based method allowed only limited powder sampling, whereas the grease-based Terasaki setup enabled multi-well sample delivery but produced higher background and weaker diffraction profiles. Conversely, Kapton-sealed Terasaki ensured secure retention of small amounts powder while providing lower background and clearer diffraction patterns. Within short total data collection times of only 1–2 min, the Kapton-Terasaki method delivered the best overall PXRD performance among the tested methods. Search-match and profile-fitting analyses showed that all three approaches sampled the same crystalline material, while the Kapton-based method gave the lowest profile residual (Rp = 9.6%) and the most reliable whole-pattern profile. These results demonstrate that optimizing sample delivery, rather than modifying the core instrument hardware, can substantially extend PXRD capability on an existing in situ crystallography platform for rapid, laboratory-based screening and comparative multi-sample measurements.

## 1. Introduction

Powder X-ray diffraction (PXRD) is one of the most widely used methods for characterizing crystalline materials, providing rapid, informative access to phase identity, crystallinity, and overall structural features without the need for single-crystal diffraction [1], [2]. For molecular and pharmaceutical powders in particular, PXRD plays an important role in screening, phase comparison, and verifying sample content [1]. At the same time, the analytical value of PXRD depends strongly on the quality of the collected diffraction signal, which is affected not only by instrumental setup but also by sample preparation, particle statistics, sample geometry, and background contributions [3]. The measured powder profile shows both the natural scattering of the material and factors from the experiment that affect the shape of peaks, how intense they are, and the background level. This makes how we handle the material an important part of the PXRD method. These factors become especially important when powder measurements are performed on instruments and pipelines originally optimized for single-crystal diffraction rather than for routine powder analysis [4]. Recent advances in laboratory X-ray instrumentation have expanded the practical flexibility of home-source diffractometers and created new opportunities to adapt available crystallographic platforms to non-standard measurement strategies [5]. In Türkiye, the installation of a modern home X-ray source in 2022 provided a new local framework for laboratory-based crystallographic data collection and method development [4], [5]. Accordingly, the Rigaku XtaLAB Synergy system allows data collection not only through the standard loop-/pin-based single-crystal pipeline, but also via the XtalCheck-S plate-reader module, primarily intended for in-situ screening of multiple crystal samples [5] (Rigaku Oxford Diffraction, 2021). Although this capability offers a practical starting point for high-throughput analysis, its direct application to powder X-ray diffraction is not straightforward, because powder specimens require a different sample-delivery logic than isolated single crystals and are more sensitive to issues such as low sample volume, poor particle statistics, and background scattering [6]. As such, the main challenge is not simply data collection itself but obtaining a powder sample geometry that yields sufficiently representative and low-background diffraction data under the limitations of a single-crystal measurement platform. To address related practical limitations in crystallographic screening, we previously developed a 3D-printed adapter that enabled Terasaki plates to be used in the XtalCheck-S module for crystal-handling applications [5]. Similarly, the present work explores whether the same general platform can be extended to powder X-ray diffraction by redesigning the sample-delivery format. Given this, Terasaki plates are cost-efficient and very useful, as they allow multi-well loading, enabling data collection from multiple samples at a time [7], [8]. However, successful practice needs a support strategy that can keep very small amounts of powder, remain compatible with the plate-reader geometry, and minimize undesirable background noise. This need is consistent with broader diffraction studies showing that thin, X-ray-transparent support materials and low-scattering sample-delivery method can substantially improve measured data quality by reducing parasitic background contributions [9], [10]. Here, we developed a modified Terasaki-plate-based sample-delivery method for PXRD on a laboratory single-crystal diffractometer implemented with the XtalCheck-S module. The method was evaluated by comparing a standard loop/pin-based method with a grease-based Terasaki strategy. Attention was focused on diffraction signal quality, practical handling, multi-well data collection, and the reproducibility of the resulting powder diffraction after batch-mode data processing. Using 5-{[4-(2-Methoxyphenyl)piperazin-1-yl]methyl}-4-ethyl-4H-1,2,4-triazole-3-thiol as a model analyte [11], we noticed that modification of the Terasaki plate with Kapton-sealed wells provides a simple and low-cost method to obtain clearer and more representative PXRD data than can be collected with either loop-based or grease-based methods under the present experimental conditions. Beyond the specific analyte examined here, the study aims to demonstrate that improving sample-delivery can largely extend the PXRD use of an available in-situ crystallography platform.

## 2. Materials and methods

### 2.1. Materials

The powder sample examined in this present study corresponds to 5-{[4-(2-Methoxyphenyl)piperazin-1-yl]methyl}-4-ethyl-4H-1,2,4-triazole-3-thiol, previously reported as compound 3 in the synthesis study of highly substituted piperazine–azole hybrids by Mermer et al [11]. This molecule contains a 1,2,4-triazole-3-thiol core linked to a 2-methoxyphenyl-substituted piperazine moiety through a methylene bridge, thus combining two pharmaceutically relevant heterocyclic fragments within a single scaffold. In the original report, the compound has been prepared from the corresponding hydrazide precursor, characterized by elemental and spectroscopic analyses (NMR, MS, etc.), and further identified as a biologically active intermediate for subsequent derivatization. Compound 3 showed measurable antimicrobial activity against both Gram-negative and Gram-positive bacteria, as well as Mycobacterium smegmatis, supporting its relevance as a structurally defined and chemically meaningful model analyte for the present PXRD method-development study.

### 2.2. Methods

#### 2.2.1 Instrumentation

Powder diffraction experiments were performed at Turkish Light Source using a Rigaku XtaLAB Synergy diffractometer equipped with a HyPix-Arc 150° detector and the XtalCheck-S plate-reader module. The instrument was used in two different operational modes: ***(i)*** the standard loop/pin-based single-crystal mounting setup [4] and ***(ii)*** the plate-reader mode enabled by our previously developed 3D-printed adapter compatible with Terasaki plates [5].

#### 2.2.2 Standard loop/pin-based powder-delivery

For the standard powder-delivery method, the powder sample was first spread on a glass slide and then looked under a microscope to identify a suitable amount for manual picking. The selected powder was then delivered using a sample pin coated with Shin-Etsu HIVAC-G grease, which was also used for powder delivery and pin mounting. Sample preparation was performed by standard laboratory tools, including a spatula, tweezers, glass slides, and a mortar and pestle, together with a tungsten needle probe, a magnetic wand, a sample pin, and a sample puck for mounting and handling. The mounted loop or pin was placed on the Intelligent Goniometer Head (IGH) of the diffractometer, and PXRD data were collected within 1 m 40 s. The detector-to-source distance was set to 120 mm, the resolution limit to 2θ ≤ 50°, and the exposure time to 15 s. The difference between the cumulative exposure time (∼75 s) and the total experimental duration reflects additional instrumental overhead, including detector readout and goniometer movements. To minimize bias arising from differences in angular coverage between the loop-based and Terasaki-based methods, the comparison emphasizes peak position consistency and relative profile quality rather than absolute intensity distributions over the full 2θ range.

#### 2.2.3 Grease-Terasaki-based powder-delivery

To adapt the plate-reader module for powder screening, powder samples were loaded into unmodified Terasaki plate wells containing grease. Grease was used to keep the powder within the wells during vertical positioning in the XtalCheck-S module, thereby enabling PXRD data collection in plate-reader mode. Data-collection parameters were optimized to improve both angular coverage and data-collection efficacy. The detector distance was increased to 120 mm, and the 2θ range was extended from 0° to 126°. In addition, omega scans were performed with 0.2° increments over the accessible scan range, and the exposure time was reduced to 0.2 s per frame. Up to 42 frames were collected for each of three independent replicates (total of 126 frames), enabling batch-mode merging after data collection within 1 m 40 s. The total effective exposure time corresponds to approximately 25 s; however, the overall experimental time (1 min 40 s) includes additional instrumental overhead, such as detector readout, goniometer motion between frames, and plate positioning. Because the grease-based setup introduced additional non-crystalline noise, it gave mainly a diffraction background, reducing overall profile quality. Overall, this method provided a cross-check for the standard loop-based method vs. Kapton-Terasaki method that is mentioned the following step.

#### 2.2.4 Kapton-Terasaki-based powder-delivery

For the optimized sample-delivery strategy, the bottom of each Terasaki well was perforated and sealed with Kapton tape (polyimide). Small amounts of powder were then loaded directly onto the Kapton-supported wells. The modified Terasaki plate was integrated to our previously developed 3D-printed plate-holder adapter [5], which enabled stable mounting and compatibility with the XtalCheck-S module. This design was aimed at minimizing sample use, reducing background noise, and supporting multi-well PXRD measurements under plate-reader geometry. After loading, the plate–adapter assembly was introduced into the XtalCheck-S module for data collection. The detector distance was arranged to 120 mm, and the 2θ range was extended from 0° to 126°. In addition, omega scans were performed with 0.2° increments over the accessible scan range, and the exposure time was reduced to 0.2 s per frame. 42 frames were collected per measurement, with an exposure time of 0.2 s per frame, yielding an effective acquisition time of ∼8.4 s per replicate. For batch-mode processing, three replicate datasets were collected (total of 126 frames), yielding a cumulative exposure time of ∼25 s. The total experimental duration (∼1 min 40 s) reflects additional overhead associated with detector readout, ω-step movements, and plate positioning within the measurement cycle. Representative optical views recorded at approximately ω = −17°, −1°, and +30° are shown in Figure 5, corresponding to the beginning, middle, and end of the omega oscillation range.

#### 2.2.5 Data processing

After data collection, diffraction frames were processed in CrysAlisPro. Processing was initiated from the main window, and the export/reprocessing settings were defined in the inset window; here, the output format was selected as HighScore (*.asc), with the exposure-time rescaling correction, smoothing, and baseline-correction options, and the calibration-information setting. The resulting 2D diffraction images were then calculated to generate 1D powder diffraction profiles (intensity vs. 2θ). For the Terasaki-based datasets, frame merging was performed in batch mode to improve particle statistics and obtain more representative powder patterns from multiple-order measurements (42 x 3). Namely, the batch mode option was selected first, followed by selecting the relevant raw diffraction files for processing. The selected settings were then confirmed by clicking OK, and the batch reprocessing/export step was run by clicking the final processing command to generate the merged powder diffraction output.

#### 2.2.6. Search–match and profile-fitting analyses

Processed powder diffraction profiles were further examined using *Match! 4* software to assess phase assignment and compare diffraction quality across the three sample-loading strategies. Search–match analysis was performed against the Crystallography Open Database (COD) under inorganic/organic-compound restrictions first. Elemental and compositional filters were applied based on the model analyte, including its empirical formula and molecular-ion information (m/z = 307.36, M+1). Following the standard *Match!* workflow, the diffraction data (*.asc) were imported, the processed patterns were refined for peak detection, and search–match calculations were then performed against the selected database. Because the analyte examined in this study is a newly synthesized organic compound that is not represented by an appropriate reference entry in the database, no meaningful phase match was obtained; so that, the search–match procedure was used primarily to evaluate database compatibility and to examine the consistency of peak positions among the loop-based, grease-based, and Kapton-based datasets under the compound restrictions alone. To compare diffraction quality more directly, profile-fitting analysis was performed on the three experimental PXRD datasets. The quality of fit was evaluated using the profile residual (Rp), calculated from the difference between observed and calculated intensities across the full diffraction profile. Accordingly, Rp values, peak-position agreement, signal intensity, and background level were compared to determine which sample-delivery method provided the most reliable representation of the same crystalline material under the present experimental conditions.

## 3. Results

### 3.1. Standard single-measurement-loop/pin method and its limitations

PXRD data were collected in standard single-crystal operational mode and sample-mounting procedures. For sample preparation, the powder was first spread on a glass slide (**Fig. 1a**) and examined under a microscope (**Fig. 1b-c**); the most suitable particles were then picked up using a standard loop (**Fig. 1d**) or a pin coated with Shin-Etsu HIVAC-G grease (**Table 1**).

**Figure 1.**
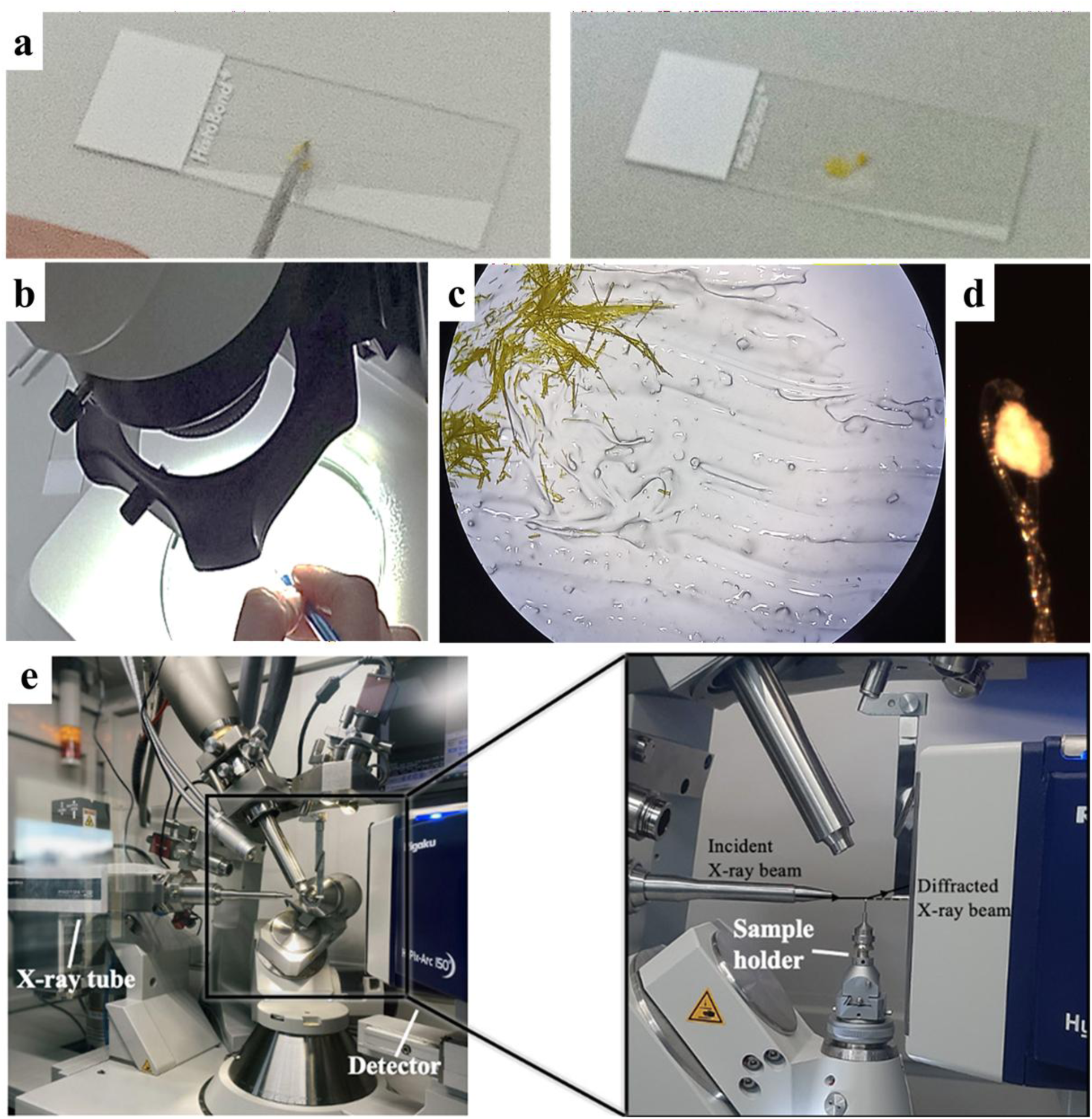
Standard loop-based powder sample preparation and mounting procedure for PXRD data collection. (a) A powder sample was spread on a glass slide, and selected particles were collected using a grease-coated loop. (b) Microscopic inspection during sample picking. (c) Representative microscopic view of the powder dispersed on the glass slide. (d) Powder is mounted on the loop prior to measurement. (e) Experimental setup on the Rigaku XtaLAB Synergy diffractometer, showing the X-ray tube, detector, and mounted sample. The magnified inset highlights the sample position relative to the incident and diffracted X-ray beams (Panel e, adopted from Atalay et al. [4]).

**Table 1.**
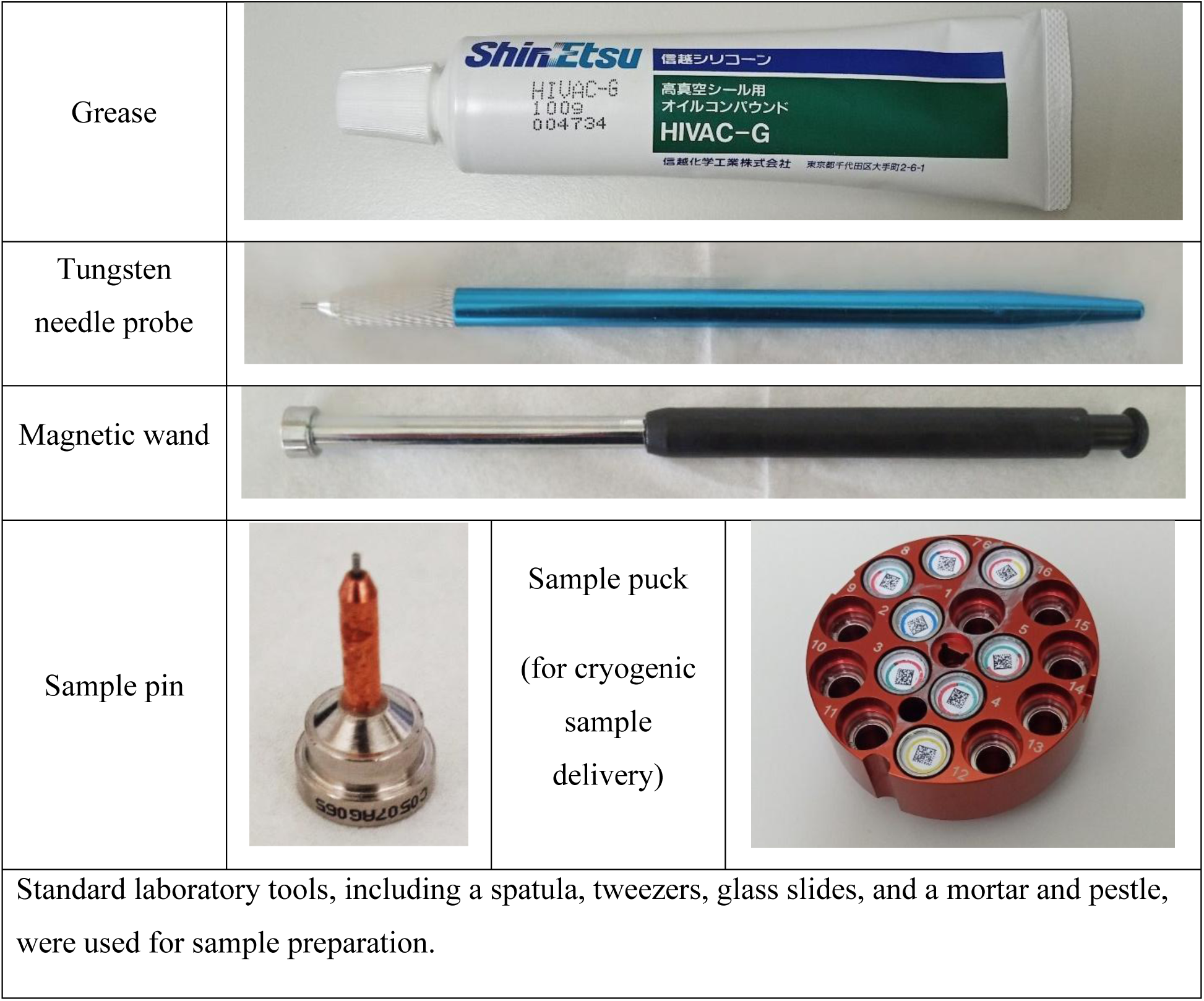
Sample-delivery tools for powder X-ray diffraction.

|  |  |  |
| --- | --- | --- |
| Grease |  |  |
| Tungsten needle probe |  |  |
| Magnetic wand |  |  |
| Sample pin |  | Sample puck<br>(for cryogenic<br>sample<br>delivery) |
| Standard laboratory tools, including a spatula, tweezers, glass slides, and a mortar and pestle, were used for sample preparation. |  |  |

The mounted pin was placed on the Intelligent Goniometer Head (IGH) of the Rigaku XtaLAB Synergy diffractometer for data collection (**Fig. 1e**) [4]. The parameters of standard powder data collection were kept as close as possible to those used in Terasaki modified methods to maintain comparable measurement conditions with a similar data collection time (1 m 40 s) while acknowledging differences in angular coverage. Accordingly, the HyPix-Arc 150° detector-to-source distance was set to 120 mm, the resolution limit to 2θ ≤ 50°, and the exposure time to 15 s, such that only five frames could be collected without exceeding the target total data-collection time of 1 min 40 s. Following data collection, the diffraction images were processed to assess the quality of the powder signal. Representative 2D diffraction frames and the corresponding integrated powder pattern obtained from these images are shown in **Figure 2**. Although this method enabled initial PXRD measurements, it demonstrated clear limitations when applied to powder samples. In practice, the loop-based setup yielded only a few frames, the diffracted intensity remained low, and the amount of powder that could be mounted remained limited. Because reliable powder diffraction data need a necessary large number of randomly oriented crystallites within the X-ray beam, these conditions were not enough to provide robust particle statistics [3], [6]. Consequently, the standard loop-based method was enough for preliminary screening within the allotted measurement time, but it was less suitable for standard PXRD measurements that need more representative and statistically robust data. Thus, the principal limitation was not the feasibility of collecting PXRD data itself, but the limited statistical quality of the data obtained within the fixed 1 min 40 s collection time. These limitations motivated the development of an alternative sample-delivery method capable of accommodating more measurements, improving signal quality, and enabling more efficient batch data collection [5], [12], [13] within a few minutes.

**Figure 2.**
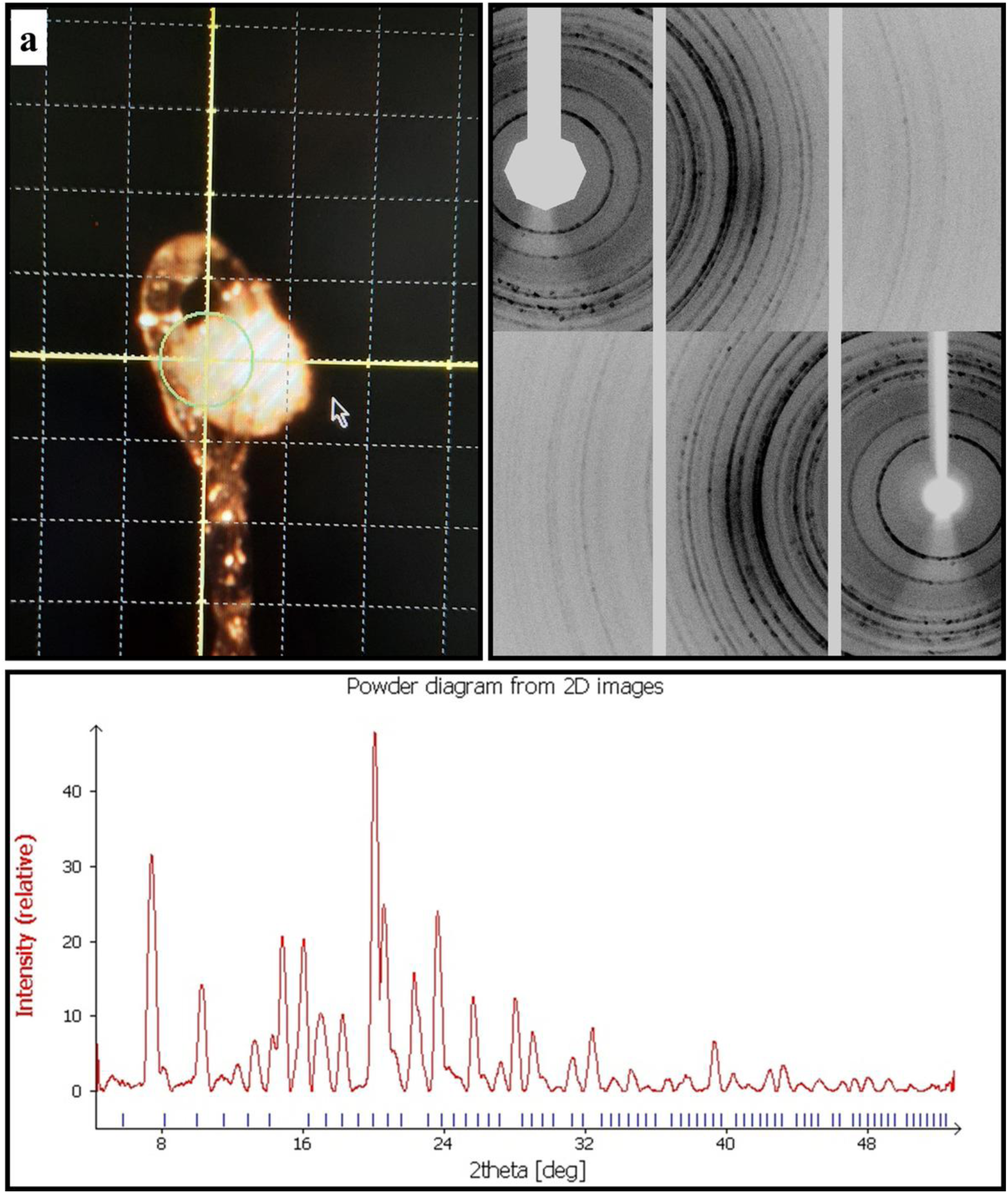
Representative diffraction obtained from the standard loop-based PXRD measurement after data collection and initial processing. (a) Optical view of the powder sample mounted on the loop and centered in the X-ray beam. The near panels show representative 2D diffraction images collected from the mounted sample, displaying powder diffraction rings. (b) One-dimensional powder diffraction pattern generated by integration of the 2D diffraction images, shown as intensity versus 2θ.

### 3.2. High-throughput Terasaki plate modification: grease– and Kapton-based approaches

To overcome the limitations of the standard loop-based method regarding both data quality and throughput, the XtalCheck-S plate reader mode, originally designed for in situ multiple-crystal screening [5], was adapted for powder X-ray diffraction measurements. In this method, 72-well Terasaki plates could be used to enable batch data collection from multiple samples at a time [5]. Here, powder samples were loaded directly into Terasaki wells containing grease, without any physical modification of the plate (**Fig. 3a**). Although this method enabled data collection, it imposed several practical limitations. In particular, the sample volume could not be reduced enough, and the delivery method was not ergonomically convenient. In addition, as also noted in the literature, amorphous binders such as grease or petroleum jelly may reduce data quality by absorbing X-rays and/or increasing background noise [12]. Consistently, the diffraction data collected using the grease-based method showed a well-defined, but relatively weaker, diffuse signal (**Fig. 3b-c**). To address these limitations, an alternative and more effective modification of the Terasaki plate was developed. The bottom of each well was perforated and sealed with Kapton tape (polyimide), a material known for its high X-ray transparency (**Fig. 4a**) [14], [15]. This modification enabled loading powder samples into the wells in very small amounts while preventing sample loss. Polymer films such as Kapton and Mylar are widely recognized as suitable supports in transmission and diffraction experiments because of their low mass absorption coefficients and minimal interference with diffraction signals [14]. In line with these properties, the Kapton-based setup provided a stronger, clearer diffraction signal (**Fig. 4b-c**) than the grease-based method (**Fig. 3b-c**), while also offering a more practical pipeline for rapid, high-throughput PXRD data collection (**Fig. 5-6**) within a few minutes alone.

**Figure 3.**
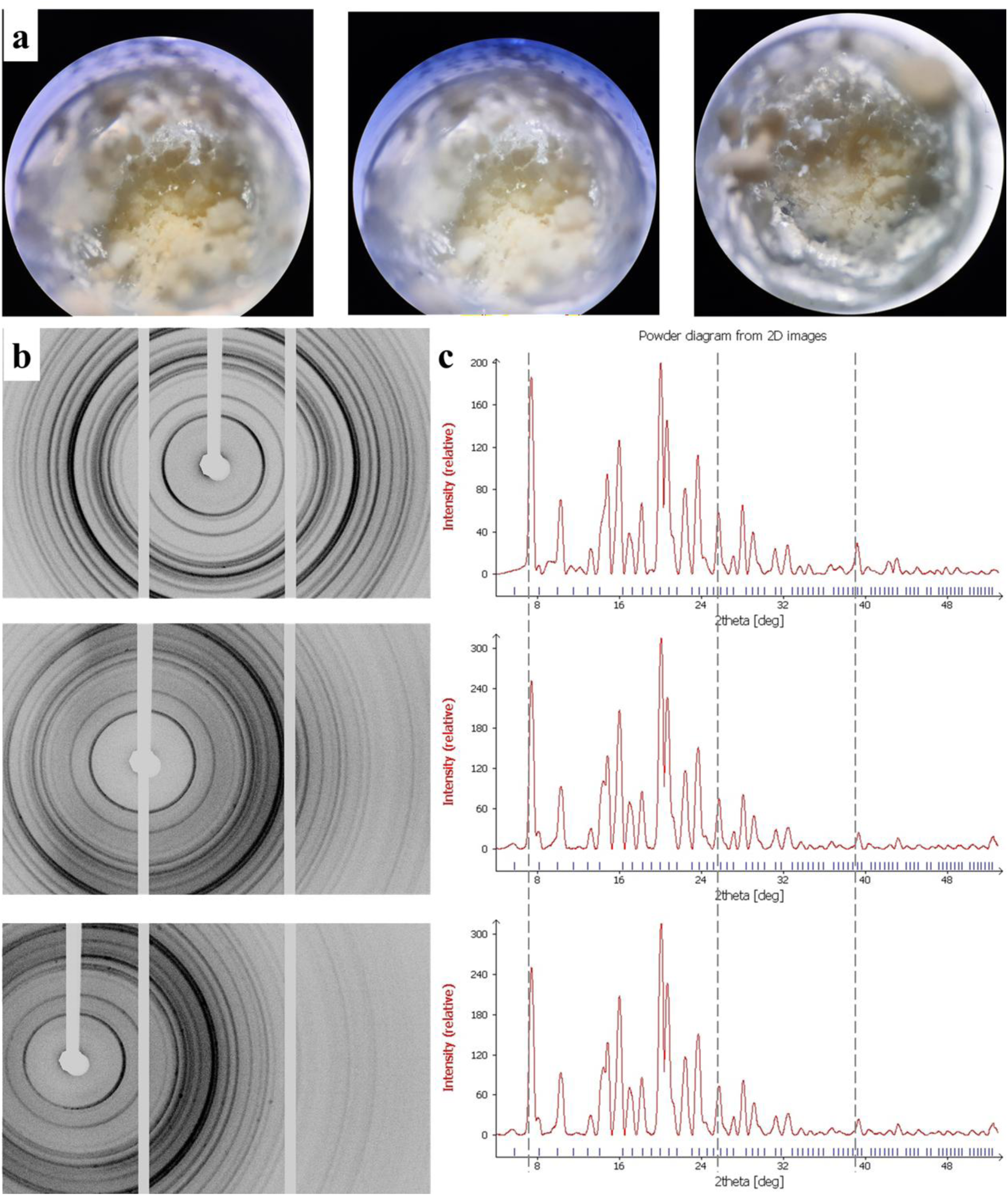
Powder X-ray diffraction data collected from grease-loaded Terasaki wells. (a) Representative optical images of powder samples placed in grease within Terasaki plate wells. (b) 2D diffraction frames obtained from the grease-based sample-delivery setup. (c) Corresponding integrated powder diffraction profiles derived from the 2D images. For batch-mode processing, 42 frames × 3 replicate wells were processed, totaling 126 frames within 1 m 40 s.

**Figure 4.**
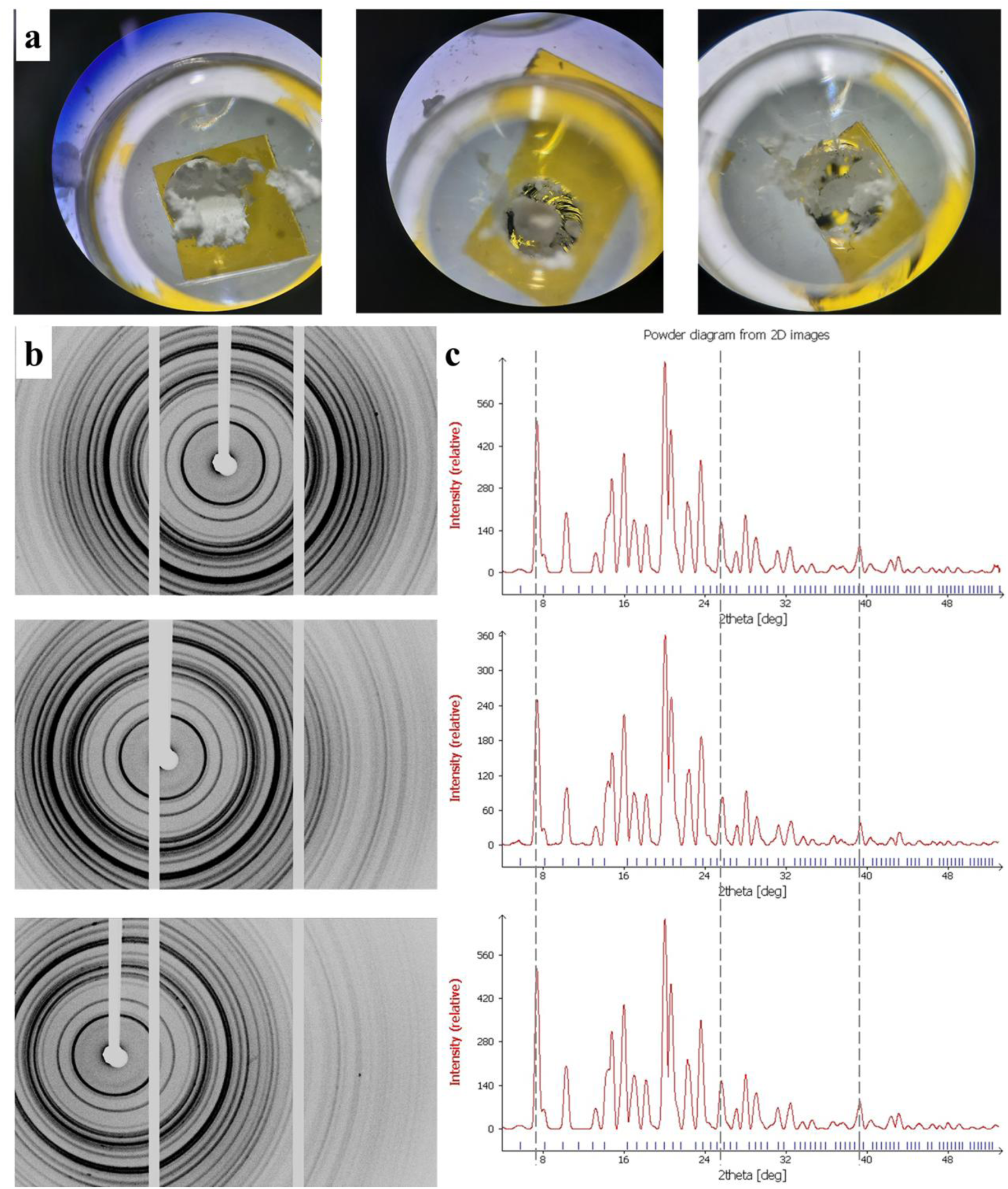
Kapton-Terasaki plate model for powder X-ray data collection. (a) Representative optical images of powder samples loaded into modified Terasaki wells sealed with Kapton tape. (b) Corresponding 2D diffraction images collected from the Kapton-supported powder samples, showing well-defined powder diffraction rings. (c) Integrated 1D powder diffraction patterns generated from the 2D images. For batch-mode processing, 42 frames × 3 replicate wells were processed, totaling 126 frames within 1 m 40 s.

**Figure 5.**
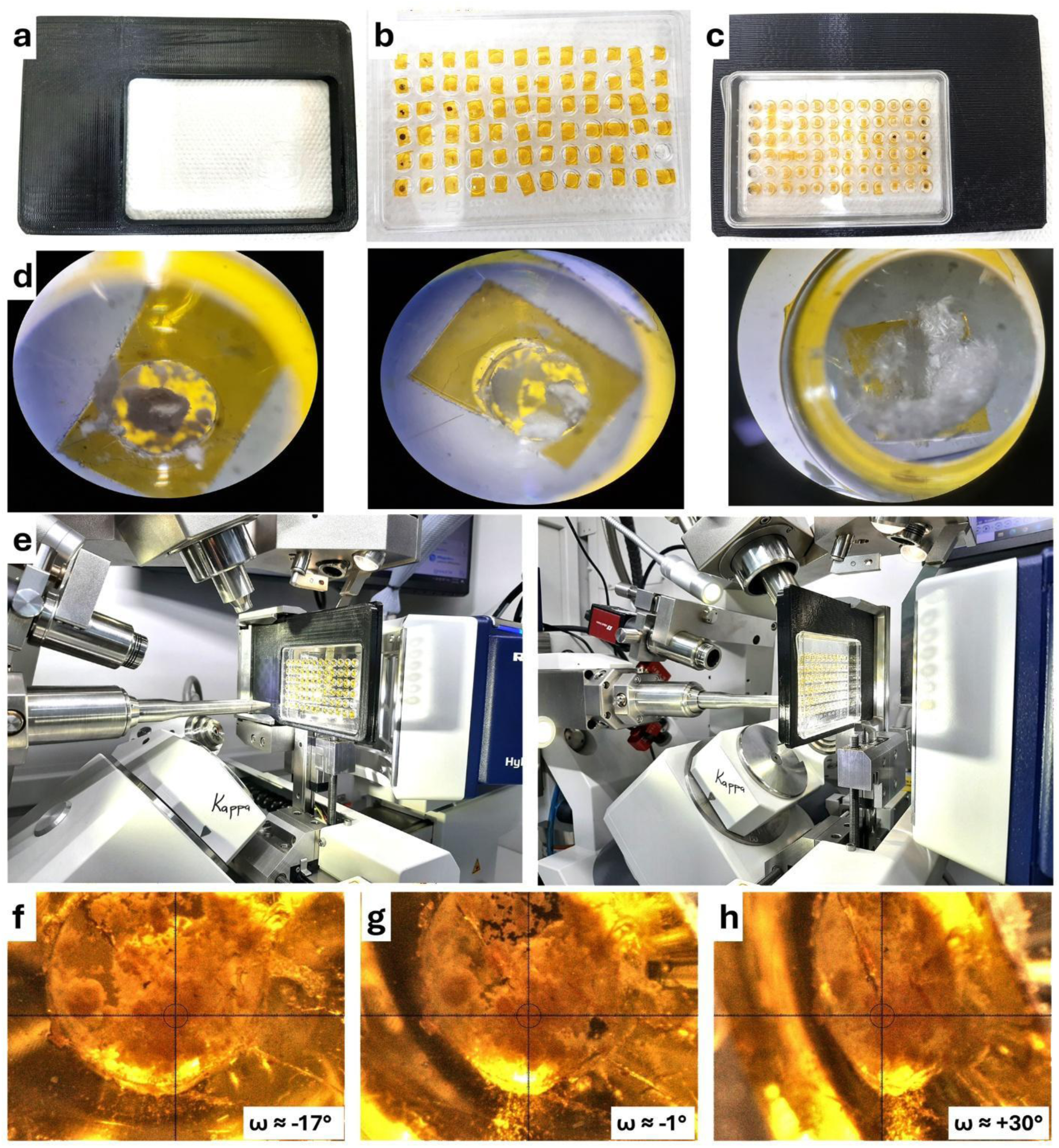
Kapton-Terasaki plate setup for high-throughput PXRD data collection using the XtalCheck-S module. (a) Previously developed 3D-printed plate-holder adapter [5]. (b,c) Modified Terasaki plate prepared for insertion into the adapter and compatible with the plate-holder adapter. (d) Representative powder samples loaded in small amounts onto Kapton-sealed wells. (e) Experimental setup showing the mounted plate–adapter assembly positioned in the XtalCheck-S module for PXRD data collection. (f–h) Representative optical views of the sample at approximately ω = −17°, −1°, and +30°, corresponding to different stages of the ω oscillation during data collection.

**Figure 6.**
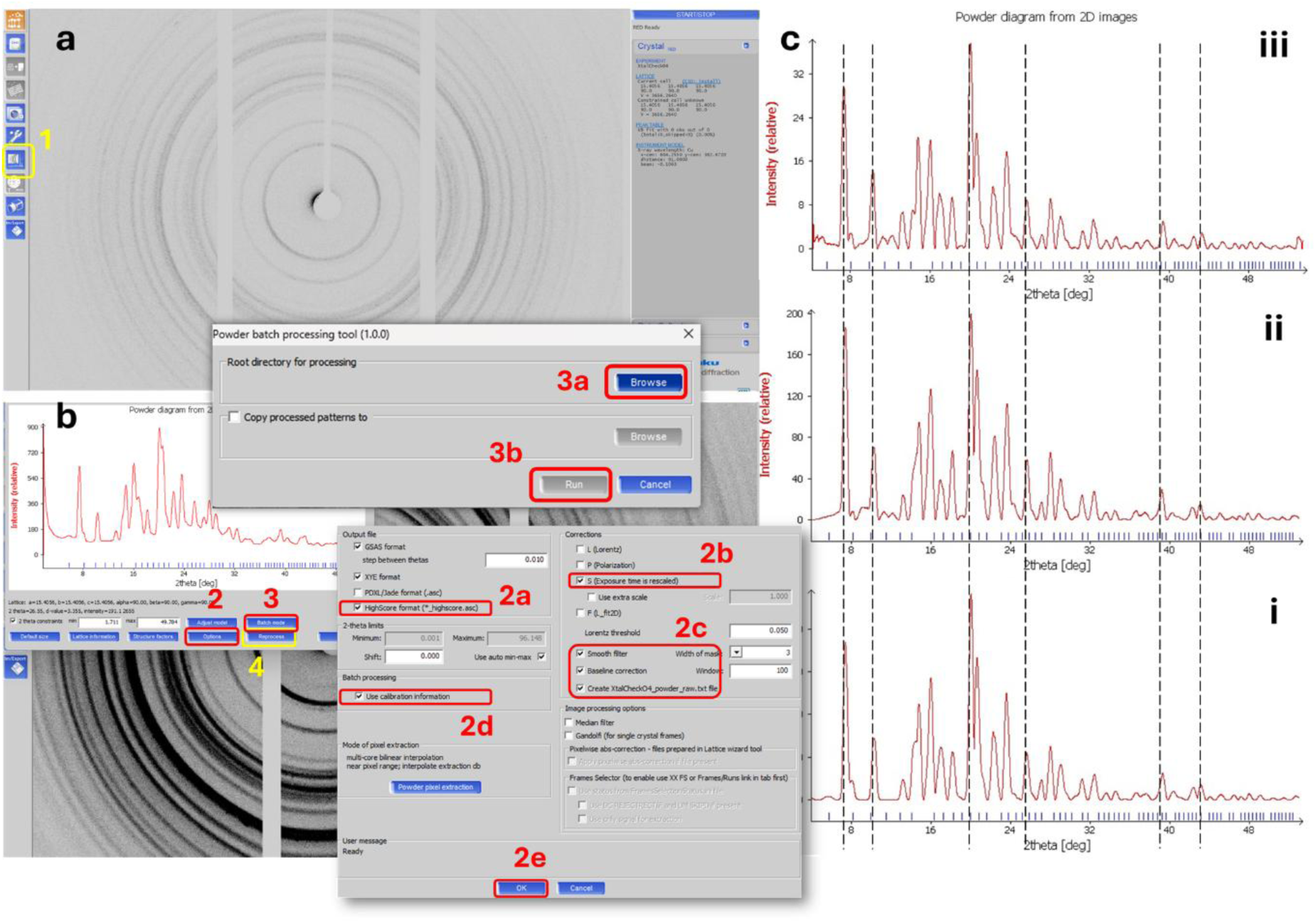
Data processing pipeline in CrysAlis^Pro^ and comparison of powder diffraction profiles obtained using the three sample-delivery methods. (a) Representative 2D diffraction file opened in CrysAlis^Pro^; the yellow box (#1) indicates the processing/reprocessing command used to initiate data treatment after collection. (b) Export and reprocessing window used to (#2) generate the final powder diagram from the 2D images. In this panel, #2a indicates selection of the output format as HighScore (*.asc), #2b indicates the selection of the exposure-time rescaling correction, #2c indicates application of smoothing and baseline-correction settings, and #2d indicates the selection of the calibration-information option in the processing settings. The numbered labels highlight the processing sequence: #3, selection of the batch raw-data via #3a brows followed by #3b run; and #4, execution of the reprocessing/export step to produce the final diffraction profile. (c) Comparison of the integrated 1D powder diffraction patterns obtained from the three sample-delivery strategies: iii, standard loop-based collection; ii, grease-based Terasaki well collection; and i, Kapton-based modified Terasaki well collection. The vertical dashed lines mark corresponding peak positions across the three datasets, showing that all methods produced similar diffraction peaks, while the Kapton-based approach yielded the strongest overall intensities (∼700 a.u.).

Accordingly, once the modified Kapton-Terasaki plate that is loaded with an appropriate amount of powder sample is installed into the 3D-printed plate-holder adapter (**Fig. 5a-d**), the assembly is introduced into the XtalCheck-S module for PXRD data collection (**Fig. 5e**). The parameters were optimized to improve both angular coverage and data-collection efficacy. The detector distance was increased to 120 mm, a scan width of 0.20° was used, and the exposure time was reduced to 0.2 s per frame. Under these conditions, 126 frames were collected in a total time of 1 m 40 s. Representative diffraction images collected at approximately ω = −17°, −1°, and +30° are shown to illustrate the beginning, middle, and end of the oscillation range (**Fig. 5f-h**).

After multi-well data collection, the diffraction frames were reprocessed in CrysAlis^Pro^ (**Fig. 6a**) using the HighScore *.asc output format together with smoothing and baseline-correction options (**Fig. 6b**). This pipeline enabled the collection of 126 frames in total, followed by merging in batch-mode within CrysAlis^Pro^. The combination of many frames with replicate-based merging improved particle statistics by increasing the representation of randomly oriented crystallites in the X-ray beam. As a result, the modified Kapton-Terasaki plate provided more robust results (**Fig. 6ci**) and an efficient PXRD pipeline (**Fig. 5**) than the grease-based (**Fig. 3** and **Fig. 6cii**) and standard loop-based methods (**Fig. 1** and **Fig. 6ciii**), while also being well suited for rapid multi-sample analysis.

To better separate differences in diffraction quality from possible differences in phase identity, the processed powder profiles were further examined by database-assisted search–match and profile-fitting analyses. Search–match analysis was performed in *Match!4* software using the COD database with organic-phase restrictions. As expected for a newly synthesized organic compound [11], no satisfactory database match was obtained for the analyte investigated here (**Fig. 7**). This result is consistent with the fact that the powder sample corresponds to the methoxyphenylpiperazine–triazole derivative reported as compound 3 in the original synthesis study [11] and is therefore not represented by an appropriate reference powder entry in the database used here. Importantly, the absence of a database match did not indicate any inconsistency among the three PXRD datasets (**Fig. 8**). On the contrary, the profiles obtained from the loop-, grease-, and Kapton-based setup showed the same characteristic reflection positions, indicating that all three methods probed the same crystalline material. To compare the quality of the measured profiles more directly, profile-fitting analysis was performed on all three experimental datasets. In each case, the overall diffraction pattern was reproduced satisfactorily, further supporting phase consistency across the powder-delivery strategies. The quality of the fit was evaluated using the profile residual (Rp) [1], defined as 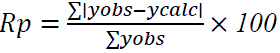 which quantifies the absolute difference between observed and calculated intensities across the diffraction profile [16]. Regarding our analysis, the fitting residuals differed among the methods: the Kapton-based configuration yielded the lowest Rp value (9.6%), followed by the loop-based measurement (12.5%), whereas the grease-based approach gave the highest residual (21.2%). Together with the stronger diffraction signal and lower background contribution observed for the Kapton-Terasaki measurements, these findings indicate that the Kapton-Terasaki setup provides the most reliable representation of the powder sample under the present experimental conditions.

**Figure 7.**
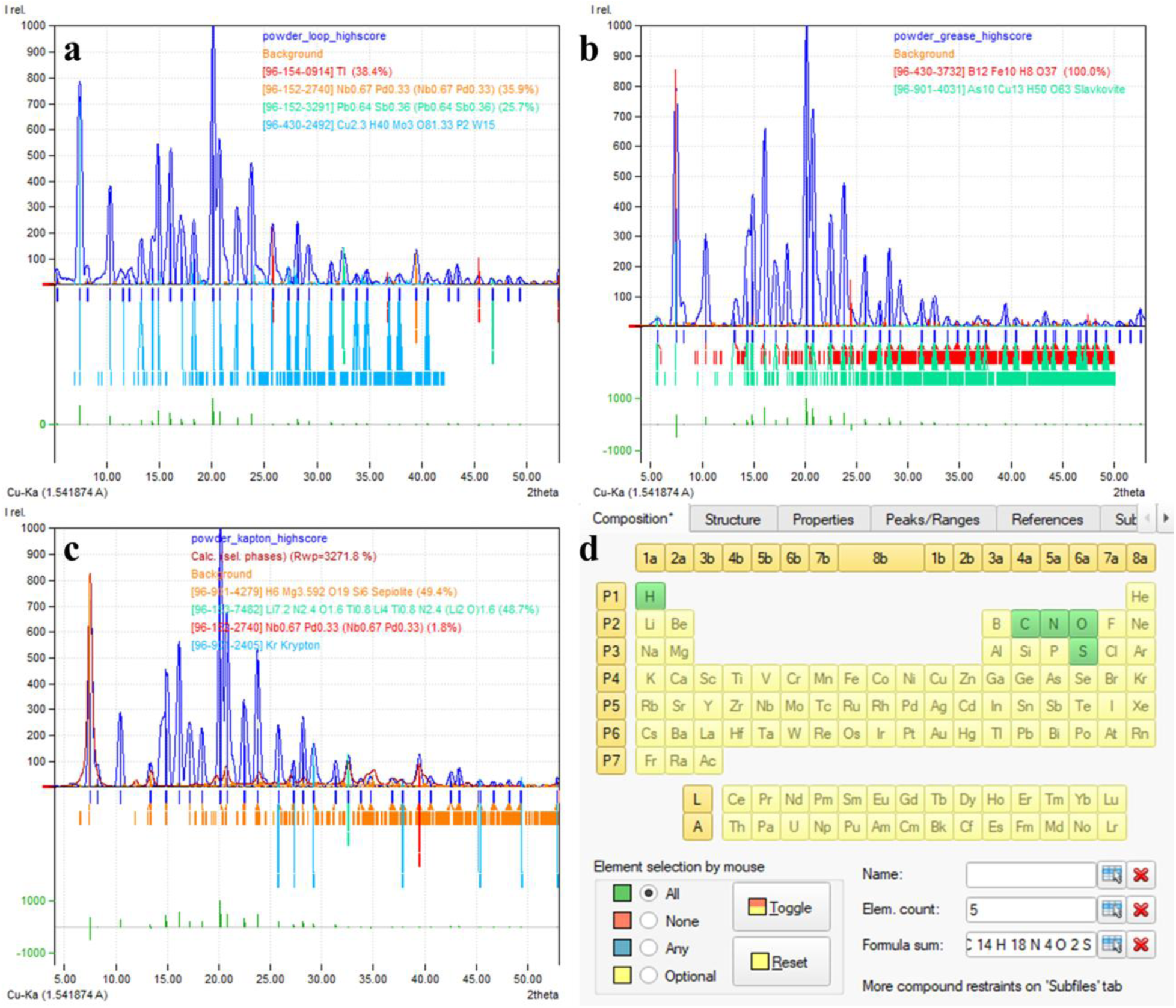
Database-assisted search–match analysis of PXRD data obtained from different sample-delivery methods. (a) Search–match results for the loop-based measurement. (b) Search–match results for the grease-based measurement. (c) Search–match results for the Kapton-Terasaki measurement, showing no satisfactory phase assignment and high mismatch between experimental and calculated patterns. (d) Elemental and compositional parameters applied during the search–match analysis, including selection of organic compounds and specification of the empirical formula (C14H18N4O2S; m/z=307.36, M+1) [11].

**Figure 8.**
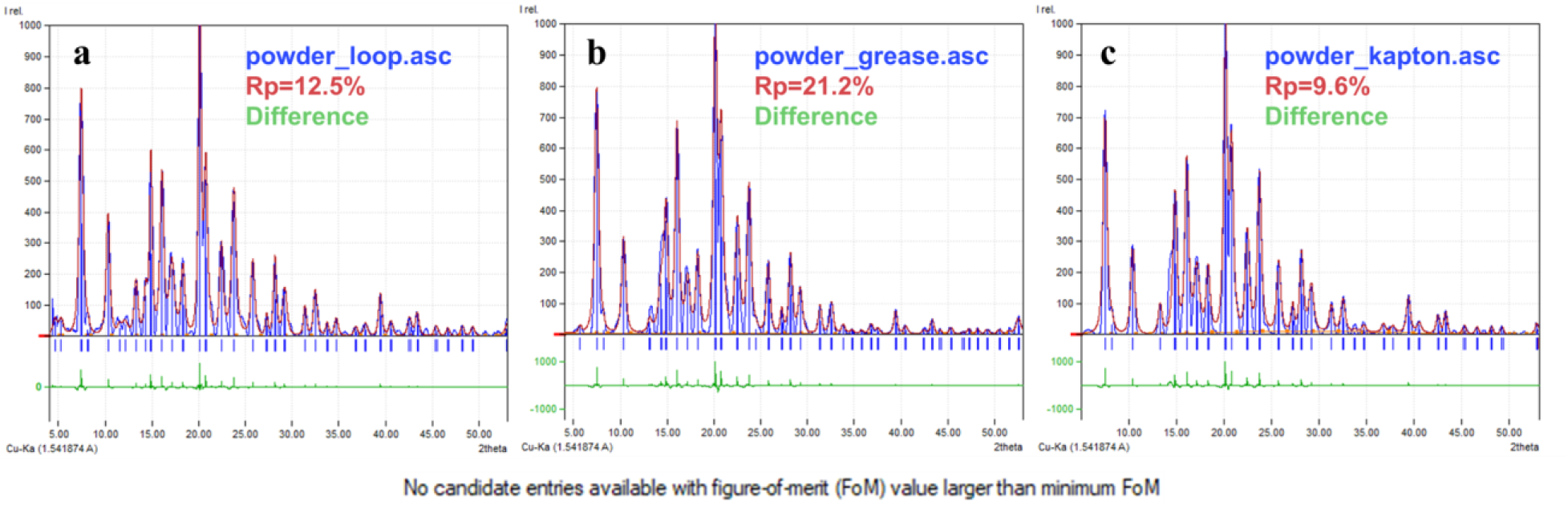
Comparison of powder diffraction profiles obtained using three different sample-delivery methods. (a) Loop-based measurement, (b) grease-based measurement, and (c) Kapton-Terasaki plate measurement. While three methods show no candidate entries available with FoM in *Match! 4*, all yield consistent peak positions, confirming that the same crystalline phase is probed, while differences in profile quality are reflected in the Rp values, with the Kapton-based method providing the lowest residual and best overall fit.

## 4. Discussion

The present study shows that a single-crystal diffractometer equipped with a plate-reader module [5] can be adapted into a practical platform for powder X-ray diffraction by redesigning the sample-delivery setup rather than modifying the core instrument hardware [4], [5]. The results indicate that data quality was determined not only by detector and scan parameters, but also by the method of introducing the powder into the X-ray beam (**Fig. 1** and **Fig. 5**). This observation is consistent with the orthodox principles of powder diffraction, in which reliable intensity data depend on sufficient particle statistics, enough sampling of randomly oriented crystallites, and the minimization of artifacts arising from specimen geometry, background scattering, and preferred orientation [17]. Accordingly, the traditional loop-based setup provided a useful starting point, but it was intrinsically limited for practical PXRD measurements, since only a very small amount of powder could be mounted, giving disposable measurements alone [12], and only a limited number of diffraction frames could be collected, whereas many frames should be collected for data merging instead. Therefore, too few crystallites contribute to the diffraction event to achieve a representative powder pattern [17].

The novel method, which moved from the loop-based format to Terasaki-plate-based method, addressed two practical issues simultaneously: throughput and diffraction accuracy. ***(i)***, the multi-well format enabled serial or batch data collection at a time (using a single Terasaki plate), offering a clear, standardized, and ergonomic setup over repeated manual pin-mounting of individual loop samples (**Fig. 5**). ***(ii)***, the Terasaki enabled evaluation of the powder in different support setups (**Fig. 3** and **Fig. 4**). The grease-containing wells showed that higher powder samples could be measured in plate-reader mode (**Fig. 3**); however, they also revealed an important limitation. Although well diffraction patterns were obtained, they were weaker, less distinct, and noisier due to the plate and grease background. This is a reasonable necessity for a bonding agent setup, as non-crystalline mounting media (such as plate and grease) may increase background and reduce the diffraction clarity [18], [19]. By contrast, the Kapton-supported setup [14], [15] provided a cleaner interface between the powder and the X-ray beam, while also allowing small amounts of powder to be retained more securely in a vertical measurement geometry. Recent studies using fixed-target diffraction have shown that thin Kapton-based membranes are very useful for delivering samples. These low-scattering, X-ray-transparent supports were chosen to reduce background noise and improve the quality of the data [9], [10] (**Fig. 4**). A particularly important aspect of the Kapton-Terasaki setup is that it improved the statistical representativeness of the powder measurement (**Fig. 6**). Namely, large numbers of frames were collected and merged across necessary and sufficient frame replicates (up to 42 frames x 3 replicate batches), thereby increasing the effective sampling of crystallites contributing to the final diffraction profile [20]. This is central to powder diffraction methodology [16], [17] because the observed intensity distribution is strongly influenced by both the number of crystallites determined and their orientation within the beam [12], [16], [17].

Here, the Kapton-Terasaki setup produced a stronger and clearer diffraction pattern, and better overall profile quality and intensity than either the grease-containing or loop-based methods. These findings suggest that the modified Terasaki setups provided a more representative powder specimen under the present experimental conditions. The lower-profile-fitting residual (Fig. 8c; Rp = 9.6%) obtained for the Kapton-adopted data further supports this idea. Thus, the main advantage of the novelty was not simply the generation of a stronger signal, but the collection of more reliable powder diffraction data.

The further search–match and profile-fitting analyses were also important for interpreting the significance of the differences observed among the three sample-delivery strategies. No satisfactory database match was obtained for the analyte, which is not unexpected for a newly synthesized organic compound lacking an appropriate reference entry [1], [2], [11], [21] (**Fig.7** and **Fig. 8**). However, the absence of a database match should not be taken to indicate a collapse of the PXRD pipeline. Importantly, the loop-, grease-, and Kapton-Terasaki datasets showed the same characteristic reflection, indicating that all three methods were in the same crystalline phase (**Fig. 6** and **Fig. 8**). Accordingly, search–match analysis primarily defined whether a known reference phase could be assigned, whereas profile fitting provided a more informative basis for comparing how faithfully each sample-delivery method reproduced the experimental diffraction pattern. This gap is momentous because, in method-development studies, alignment in peak positions across methods differs may be more informative than definitive phase identification, particularly when the material under question is novel [1], [17], [21].

Taken together, these results suggest that the modified Kapton-Terasaki setup is more than an alternative sample holder. Rather, it represents an improved PXRD sampling strategy for a laboratory single-crystal diffractometer implemented with a plate-reader module. Its principal advantages include low sample consumption, in line with rapid multi-sample collection, stronger diffraction intensity, and more reliable whole-pattern representation. These features make the method particularly significant for preliminary screening, comparative multi-powder studies at a time, and applications in which innumerable powder samples must be examined within too limited instrument time. Additionally, the method could be considered the best practical screening and data-collection approach rather than a universal substitute for dedicated high-resolution powder diffractometry in Türkiye. As widely recognized in the powder diffraction literature [17], [21], quantitative phase analysis and full structure determination remain sensitive to specimen effects, instrumental geometry, peak overlap, and the availability of suitable reference data. However, here, the Kapton-Terasaki method provides a simple, cost-efficient, and experimentally pragmatic way to extend the PXRD capabilities of an existing in situ crystallography platform.

## 5. Conclusion

This study shows that PXRD performance on a laboratory single-crystal diffractometer can be improved through optimization of the sample-delivery strategy, without modification of the core instrument hardware. Among the configurations examined, the Kapton-Terasaki setup gave the best overall powder-diffraction performance under the present experimental conditions, combining low sample consumption, practical handling, compatibility with the XtalCheck-S module, and improved data quality. Compared with both the conventional loop/pin-based method and the grease-based Terasaki configuration, the Kapton-supported method yielded higher diffraction intensity, lower background, and a more faithful whole-pattern profile after batch-mode processing. These findings indicate that successful adaptation of single-crystal-oriented platforms to powder measurements depends strongly on sample presentation and support geometry, in addition to data collection conditions. The Kapton-Terasaki method therefore offers a simple and experimentally practical means of extending PXRD capability on an existing in situ crystallography platform, particularly for rapid laboratory-based screening and comparative multi-sample measurements within a few minutes even.

## Author Contributions

The analyte was synthesized by A.M. The Kapton–Terasaki setup was installed by E.A. and A.K., and data collection, processing, and further analysis were performed by E.A. All authors have read and approved the final version of the manuscript.

## Acknowledgements

The authors gratefully acknowledge Şaban Tekin, Jakub Wojciechowski, and Ahmet Katı for their invaluable technical and administrative support in facilitating access to and use of the Turkish Light Source.

